# Improved detection of fungi and uncultivated microorganisms in soil metagenomes using a comprehensive genome database

**DOI:** 10.1101/2025.03.21.644662

**Authors:** Zoey Werbin, Ilija Dukovski, Dylan Mankel, Winston E Anthony, Daniel Segrè, Jennifer M. Bhatnagar, Marianna Felici

## Abstract

Soils harbor diverse microbial communities crucial for ecosystem functioning, but poor genomic representation of many uncultured soil microorganisms limits the utility of existing databases to address some of the most pressing questions in environmental microbiology. To address this, we developed the SoilMicrobeDB, a comprehensive, genome-based reference database to enhance metagenomic classification for soil ecosystems, with a focus on previously underrepresented fungal taxa and uncultured organisms. We evaluated the database using a large soil metagenome dataset, comparing classification rates, analyzing fungal-bacterial ratios against phospholipid fatty acid (PLFA) estimates, and validating lineage abundances with rRNA amplicon sequencing data. Mock community analysis was also conducted to test the precision of community classification and the prevalence of false positives. The SoilMicrobeDB workflow improved metagenomic read classification by over 20% and provided more accurate fungal abundance estimates, particularly for nutrient cycling groups such as ectomycorrhizal fungi. Metagenomic-derived fungal-bacterial ratios were correlated with PLFA and qPCR estimates, and lineage proportions were aligned with relative abundances estimates from rRNA amplicon sequencing. Uncultured taxa represented up to 50% of classifiable soil microbial communities in certain biomes. SoilMicrobeDB offers robust taxonomic and functional profiling of soil communities and provides a scalable and updatable tool for soil microbial ecology research. SoilMicrobeDB is accessible through an interactive platform linking genomes to environmental factors, enabling researchers to explore microbial distributions across soil conditions and potentially leading to new insights into soil ecology and management practices.

## INTRODUCTION

Soil contains most of the microbial life in the terrestrial biosphere, but poor annotations of soil genomic datasets limit our biological and ecological understanding of their diversity and activity (Anthony et al., 2024; Edwin et al., 2024; Nesme et al., 2016). The historical lag in generating whole-genome sequences for microorganisms from complex environments has led to genomic resources that lack representation of these communities (Edwin et al., 2024; Geisen et al., 2019; Nayfach et al., 2020). Furthermore, databases such as RefSeq maintained by the National Center for Biotechnology Information (NCBI) are overwhelmingly biased toward medically-relevant microbes, and the curated phylogenetic Genome Taxonomy Database (GTDB; (Parks et al., 2015) excludes all fungi, which are critical members of the soil microbiome (Baldrian, 2019; Brabcová et al., 2016; De Boer et al., 2005; Talbot et al., 2008). Taken together, these issues limit our ability to fully characterize the microbial DNA found in soil samples (Anthony et al., 2024)). Because microbial genes can predict and explain the properties and functional characteristics of complex communities (Graham et al., 2016; Karaoz & Brodie, 2022; Rocca et al., 2015; Tomasek et al., 2017; Vietorisz et al., 2024), soil ecology research would benefit from a specialized up-to-date DNA-based repository of soil microbial taxa (Choi et al., 2017; Edwin et al., 2024).

We present SoilMicrobeDB, a customizable and searchable genome-based reference for classifying unassembled microbial DNA from soil samples. This resource contains published sequenced genomes of thousands of bacterial, archaeal, and fungal taxa, as well as over 20000 novel microbial genomes assembled from short-read environmental DNA sequences (called Metagenome-Assembled-Genomes, or MAGs, Figure 1), screened for high genome completeness and minimal contamination. This meets a previously described need for an improved soil DNA reference database (Anthony et al., 2024; Choi et al., 2017; Edwin et al., 2024) and includes hundreds of genomes from the “most wanted” soil lineages such as Verrucomicrobiota and Acidobacteriota, which are suspected to play keystone roles in soil nutrient cycling but have remained largely uncharacterized (Choi et al., 2017; Kalam et al., 2020). By increasing the identifiable microbiome composition of soil samples, SoilMicrobeDB can support insights into ecological relationships between taxonomic diversity across environments. Abundances from SoilMicrobeDB can also be linked to environmental data to identify microbes with potentially unique physiological tolerances, such as fungi detected in extreme environments (Bueno De Mesquita et al., 2024); we developed an interactive web application that illustrates this use case.

**Figure 1.**
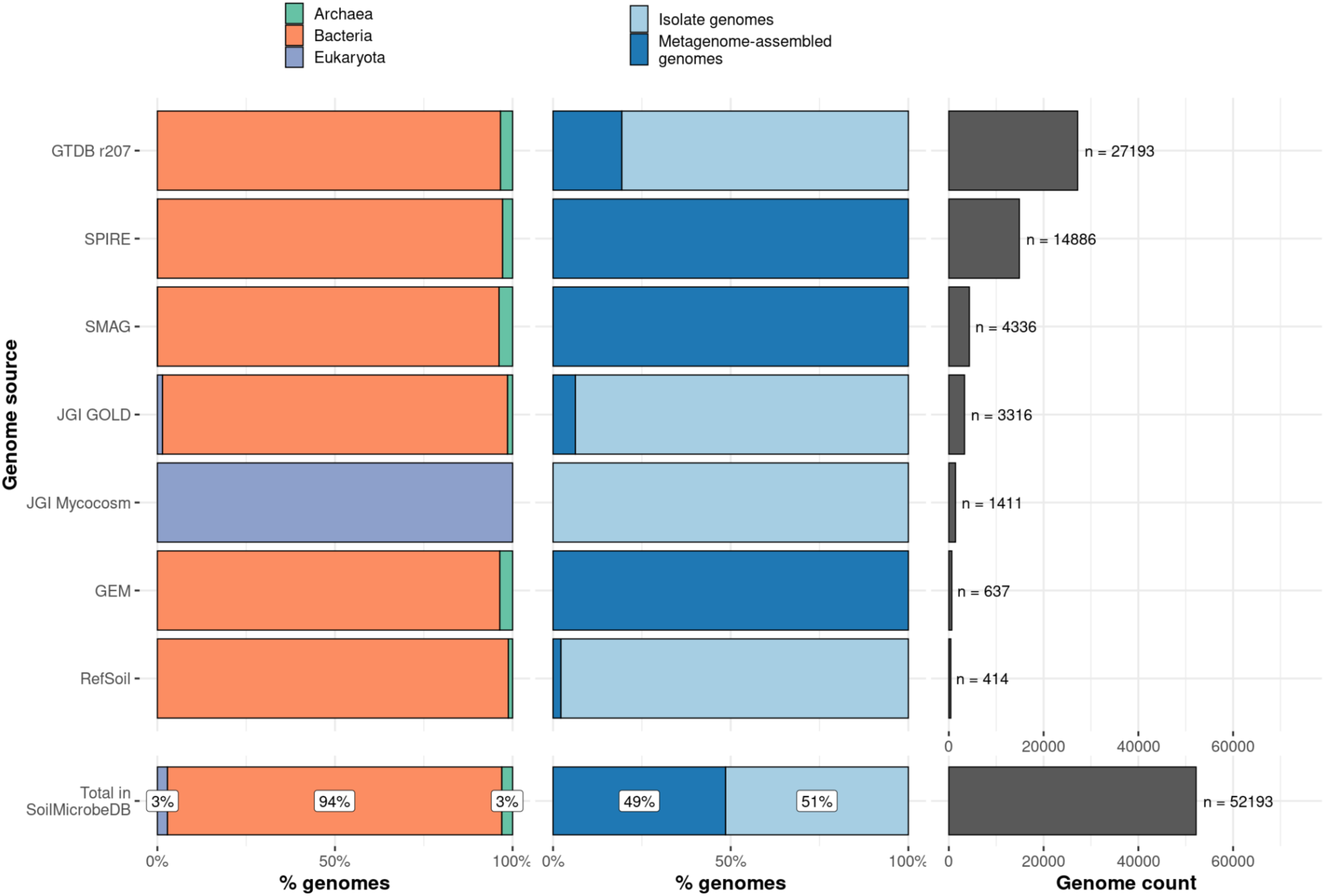
Sources of publicly available genomes included in SoilMicrobeDB. GTDB r207 genomes were filtered by 95% average nucleotide identity. MAGs were filtered by estimated genome completeness (>95%) and contamination (<5%). Left panel: proportion of genomes in major taxonomic domains. Middle: proportion of genomes derived from isolates or assembled from metagenomic reads. Right: unique number of genomes from each genome collection included within SoilMicrobeDB.

### Considerations for a comprehensive soil genome database

#### Metagenome-Assembled-Genomes

Although advances in bioinformatics have enabled the increased identification of MAGs, these assemblies are often associated with individual research projects, leaving them unlikely to be collected into a comprehensive database (Youngblut & Ley, 2021). These MAGS also frequently are missing metadata that would allow researchers to have confidence in their quality, such as contamination and completeness scores (Breitweiser et al. 2019; Bowers et al. 2017). However, recent publications of tens of thousands of soil-associated MAGs with detailed quality information and online download portals such as the Soil MAG catalog (Ma et al. 2023) and SPIRE (Searchable Planetary-scale mIcrobiome REsource; Schmidt et al. 2023) present an opportunity to better integrate MAGs into soil reference assembly databases. To facilitate the automatic creation of genome-scale metabolic models (GEMs), which can be used to simulate the activity of microbes within soil environments (Borer et al., 2019), we exclude medium- and low-quality genomes from SoilMicrobeDB. Although specific relationships between genome quality and GEM quality are not currently well-characterized (Tarzi et al., 2024), we err on the side of high-quality genomes by implementing a genome completion cutoff of 95% and a maximum contamination cutoff of 5%, resulting in 25,333 MAGs in SoilMicrobeDB. Users who prefer different cutoffs can download and customize the SoilMicrobeDB (see Appendix 1: User Guide), or separately submit unclassified sequencing reads to separate databases.

#### Fungi

Fungi are generally detected at low rates within metagenomic datasets, but this may be an issue of inadequate reference genome datasets (Donovan et al., 2018; Edwin et al., 2024). Compared to prokaryotes, there are far fewer species of soil fungi with fully sequenced genomes (65,703 prokaryotic species in GTDB, compared to 2,400 fungi in the Mycocosm collection maintained by the Joint Genome Institute). Initiatives to increase available fungal genome sequences, such as the 1000 Fungal Genomes project (Grigoriev et al., 2014), have produced high-quality genomes that have not yet been integrated into searchable genome databases. All published genomes from Mycocosm (1411) were included in SoilMicrobeDB, allowing us to leverage these initiatives to improve our annotation of fungi within soil DNA sequencing datasets.

### Benchmarking the database against other soil microbiome indicators

To demonstrate the value of the database as an investigation tool, we used SoilMicrobeDB to test the following hypotheses:

1. Expanding the soil-specific diversity of existing genome databases, in the form of novel MAGs and fungal genomes, increases the number of soil microbiome taxonomic classifications in soil metagenome data and increases the confidence in these classifications.
2. Estimates of fungal abundance obtained through metagenomics can be reconciled with estimates obtained through other methods (e.g. PLFA, qPCR, and ITS)
3. Soils with more unclassified DNA also have higher abundances of uncultured organisms (e.g. MAGs). These metagenomic properties vary across soils from different environments, indicating hotspots of under-representing microbial diversity.

To test these hypotheses, we evaluated the classification of reads from (1) simulated metagenomic communities and (2) a diverse metagenomic sequence collection of over 1400 soil samples distributed by the National Ecological Observatory Network (NEON) (Battelle Memorial Institute, 2019). Samples were classified using either existing Kraken2 databases or a custom Kraken2 genome database (the SoilMicrobeDB) created using the Struo2 pipeline (Lu & Salzberg, 2020; Wood et al., 2019; Youngblut & Ley, 2021). All classifications were filtered to high-quality assignments using kmer multiplicity and entropy (Diener et al., 2020)) and then normalized by genome length to estimate taxon abundances with Bracken (Lu et al., 2022). Species-level abundances and associated sample data can be downloaded via a new SoilMicrobeDB interactive web application. To demonstrate the utility of the dataset for identifying taxa of interest, such as extremophiles, this web application also includes exploratory visualizations showing how each microbial species is distributed across environmental gradients.

## METHODS

### Genome sources

Genomes were integrated into the SoilMicrobeDB from multiple sources (Table 1) by assigning each genome a NCBI taxonomy ID. For genomes that had been taxonomically classified using the taxonomic nomenclature of the Genome Taxonomy Database (GTDB), this required converting from GTDB to NCBI taxonomy using GTDB-provided translation files associated with each release. All MAGs had been published according to the Minimum Information about any (X) Sequence (MIxS) version 6.2.0, provided by the Genome Standards Consortium (GSC). This step allowed us to filter MAGs by quality, retaining those with over 95% completeness, less than 5% contamination, and the presence of 16S rRNA genes (Nayfach et al. 2021). MAG assemblies were cross-referenced with GTDB r207 genome accessions to prevent duplication of genomes in SoilMicrobeDB. All download scripts are available on the SoilMicrobeDB Github repository: https://github.com/zoey-rw/SoilMicrobeDB.

**Table 1.** Data sources for publicly-available genomes in the SoilMicrobeDB. MAG collections were filtered to those with over 95% completeness, less than 5% contamination, and the presence of 16S rRNA genes (Nayfach et al.. 2021). GTDB r207 was dereplicated to 95% average nucleotide identity (ANI) by Struo2 pipeline developers (Youngblut & Ley, 2021). All unpublished or restricted-access genomes were excluded.

| Source | Genome type | Domain | Count in initial collection | Count in SoilMicrobe DB | Context | Citation |
| --- | --- | --- | --- | --- | --- | --- |
| GTDB r207 (dereplicated) | Isolate and Metagenome-Assembled | Bacteria | 40456 | 26261 | Not soil-specific | (Parks et al., 2021; Youngblut & Ley, 2021) |
|  |  | Archaea | 1410 | 932 |  |  |
| JGI GOLD | Isolate | Bacteria | 7469 | 3221 | Soil-associated | (Mukherjee et al., 2021) |
|  |  | Archaea | 83 | 48 |  |  |
|  |  | Fungi | 245 | 44 |  |  |
| RefSoil | Isolate | Bacteria | 888 | 409 | Soil-associated | (Choi et al., 2017) |
|  |  | Archaea | 34 | 5 |  |  |
| Mycocosm | Isolate | Fungi | 2537 | 1411 | Not soil-specific. Low-complexity regions masked | (Grigoriev et al., 2014) |
| SMAG catalogue | Metagenome-Assembled | Bacteria | 2686 | 4169 | Soil-associated | (Ma et al., 2023) |
|  |  | Archaea | 120 | 166 |  |  |
| GEM catalog | Metagenome-Assembled | Bacteria | 339 | 614 | Soil-associated | (Nayfach et al., 2020) |
|  |  | Archaea | 5 | 23 |  |  |
| SPIRE | Metagenome-Assembled | Bacteria | 1136155 | 14453 | Soil-associated | (Schmidt et al., 2023) |
|  |  | Archaea | 21204 | 424 |  |  |

#### JGI GOLD

The Joint Genome Institute (JGI) Genomes OnLine Database (GOLD) maintains a vast collection of genomes derived from isolates and assembled from metagenomes. Soil-associated samples were identified by filtering to the presence of the term “Soil” within the “Ecosystem Type” category in GOLD’s 5-Level Ecosystem Classification Paths Excel (downloaded from JGI GOLD https://gold.jgi.doe.gov/downloads, generated 20 Jun, 2022). Organisms from these ecosystems were identified using the Organism, Sequencing, and Analysis tables within the Public Studies/Biosamples/SPs/APs/Organisms Excel (downloaded from JGI GOLD https://gold.jgi.doe.gov/downloads, generated 20 Jun, 2022). Organisms were filtered to remove any with a “Restricted” value within the “DATA UTILIZATION STATUS” column. All unique NCBI accessions were downloaded using the ncbi-genome-download utility (Blin, 2023). To decrease the bias toward medically-relevant lineages (e.g. over 600 genomes for *Streptococcus pyogenes*), a maximum of 5 randomly-selected genomes were retained per NCBI taxonomic ID, resulting in 3316 genomes.

#### GTDB r207

The Genome Taxonomy Database release 207 (GTDB r207) (Parks et al., 2020, 2021) was filtered to genomes with >50% CheckM completeness – less restrictive than our general completeness cutoff – to maintain the broad phylogenetic diversity represented by this database. As with the JGI GOLD genomes, after conversion to NCBI taxonomic IDs, a maximum of 5 randomly-selected genomes were retained per NCBI taxonomic ID, resulting in 27193 genomes from the GTDB.

#### Mycocosm

The Joint Genome Institute’s Mycocosm database (Grigoriev et al., 2014) includes a collection of fungal isolate genomes from various habitat types. All unrestricted-access fungal genomes were downloaded as repeat-masked genome files downloaded using a custom web-scraper script.

#### SMAG

This Soil Metagenome-Assembled Genome (SMAG) collection (Ma et al., 2023)included soil-associated metagenome-assembled genomes (MAGs). Metadata to filter genomes by completeness and contamination was provided in the supplement of the original manuscript. The genome names within the metadata file were used to generate download paths within R.

#### SPIRE

The Searchable Planetary-scale mIcrobiome REsource (SPIRE) collection (Schmidt et al., 2023) included environmental MAGs from various habitat types. To determine soil-associated genomes, SPIRE’s “microntology” file was filtered to rows with the following terms in the habitat column: soil, rhizo (i.e., “rhizosphere”), forest, cropland, terrestrial, litter, and agriculture. The metadata from release version 1 was then used to filter genomes by completeness and contamination. The genome names within the metadata file were used to generate download paths within R.

#### GEM

The Genomic Catalog of Earth’s Microbiomes (GEM) collection (Nayfach et al., 2020) included environmental MAGs from various habitat types, linked to models. GEM collection metadata were accessed from NERSC, and filtered to “Soil” in the “ecosystem_type” column. The genome names within the “genome_id” column of the metadata file were used to generate download paths within R.

### Viral genomes

Viruses were excluded from SoilMicrobeDB after tests against a RefSeq collection of 26,129 viral genomes showed classification rates close to zero. For viruses to be accurately detected in soil metagenomes, sample preparation would likely have to prioritize viruses at the expense of all other microbial information, using techniques like size fractionation or filtration before DNA extraction (Vibin et al. 2018; Santos-Medellin 2021). Another obstacle to combining viruses with non-viruses is presented by viral taxonomy, which is usually assigned through software that is not consistent with NCBI taxonomy. Therefore, we recommend that viruses should be classified using a separate database for specially-prepared soil samples.

### Database creation and updating

To create a Kraken2 database from locally-downloaded genomes, we used the Snakemake-based Struo2 pipeline (Youngblut et al. 2020). When provided with 28 cores, the database pipeline rns in approximately 4.5 hours and uses 37 GB of storage. We provide a User Guide (Appendix 1) with details on how to access the database through an AWS repository and update it with new genomes.

### Database evaluation with simulated mock communities

To evaluate the performance of databases using samples with known taxon abundances, metagenomic samples were generated from mock communities. We used the InSilicoSeq package in Python (Gourlé et al., 2019) to create mock communities of 200 genomes, from genera common to the SoilMicrobeDB and two commonly-used reference databases: GTDB r207, and PlusPF (Table 2). Mock communities had varying proportions (1%, 3%, 5%, 10%, 15%, and 20%) of fungal genomes, and varying numbers of simulated reads, or sequencing depths (500K, 1M, 5M) resulting in 18 mock communities. Forward and reverse read files were generated using the “generate” command with the “hiseq” flag for error profiling. Each sample was classified using Kraken2, then passed to Architeuthis for quality filtering (additional details in *Kmer-based false positive analysis*), before abundance estimation with the Bracken software.

**Table 2.** Databases used for comparison with the SoilMicrobeDB.

| Database name/version | Contents | Size (taxonomy nodes) | Size (GB) | Access |
| --- | --- | --- | --- | --- |
| PlusPF, 10-2023 | RefSeq archaea, bacteria, viral, plasmid, human, fungi, protozoa, UniVec_Core | 45903, 49820 taxonomy nodes | 74 GB | https://genome-idx.s3.amazonaws.com/kraken/k2_pluspf_20231009.tar.gz |
| Genome Taxonomy Database r207 | Archaea and bacteria from Genome Taxonomy Database r207, de-replicated to 95% average nucleotide identity | 62443 genomes, 34874 taxonomy nodes | 179 GB | Distributed for Struo pipeline: http://ftp.tue.mpg.de/ebio/projects/struo2/GTDB_release207/kraken2/ |
| SoilMicrobeDB | See Table 1 | 40364 taxonomy nodes | 280 GB | AWS FTP Link |

### Database evaluation with soil sample collection

After database creation, we evaluated the performance using 1400 samples sequenced by the National Ecological Observatory Network, representing many different soil ecosystems and a range of sequencing depths. Specific protocols are updated regularly and reported with sample metadata, with full sequencing details in the NEON Metagenomics standard operating procedure (Battelle Memorial Institute, n.d.). Briefly, double-stranded DNA was sheared using then extracted using KAPA HyperPrep Kit (Roche KAPA Biosystems, 2023). qPCR performed for quantification, followed by size selection. Samples from multiple sites are pooled into 2 sets of 96-well plates for 150 bp paired-end sequencing using Illumina NovaSeq 6000. Samples with fewer than 5 million reads are reanalyzed.

The classification step requires at least 40 GB of RAM to search against the SoilMicrobeDB, and up to 300 GB of RAM was required for the GTDB. Each sample was run against each Kraken2 database using the “report-minimizer-data” and “--report-zero-counts” flags.

### Kmer-based false positive analysis

To identify false positives, we used the Architeuthis software (Diener, 2024), which leverages a recently-developed feature of Kraken2 that reports the number of unique k-mers mapped to an individual genome. Trends in k-mer matches are informative because a genome present in the sample should have matches evenly distributed across the genome (Donovan et al. 2018), so a low value of unique k-mers may be a false positive because of low-complexity regions, misassembled or contaminated genomes, or horizontal gene transfer. Architeuthis filters out matches using three metrics for each read: multiplicity (multi-lineage matches within the same taxonomic rank), consistency (proportion of kmers with matching final classification), and entropy (multiplicity weighted by taxon abundance). Architeuthis was run on Kraken2 output with default filter parameters (--min-consistency 0.9, --max-multiplicity 2, --max-entropy 0.1). The filtered output was then passed to Bracken to estimate the abundances of each taxon within a sample.

### Fungal-bacterial ratio evaluation

To determine whether metagenome-derived fungal-bacterial ratios were consistent with estimates derived from other data sources, we leveraged various types of microbial data collected from the same soils as the metagenomic DNA extractions (described below). We also compared metagenome-derived fungal-bacterial ratios against spatially-explicit model-derived predictions from environmental variables (Yu et al., 2022). Comparisons were performed using linear regression on square-root transformed values using the “lm” function in R (R Development Core Team, 2008).

#### Quantitative PCR analysis

Quantitative polymerase chain reaction (qPCR) data were obtained from NEON DP1.10109.001 for all available sites and dates using the neonUtilities R package (Lunch CK, 2020). Detailed data collection methods are available in (L. Stanish, 2023). Soil samples were collected and frozen on dry ice before being transferred to ultra-low-temperature freezers for storage and shipment to an external laboratory for processing. qPCR was performed targeting the 16S subunit of the rRNA gene operon to estimate abundances of bacteria and archaea and the internal transcribed spacer (ITS) region of the rRNA gene operon to estimate the abundance of fungi (Stanish et al. 2023). Primers 341F 805R were used for bacteria and archaea, and ITS1f and ITS2r primers were used for fungi. Gene abundances were calculated using the mean of three replicates per DNA sample.

#### PLFA (phospholipid fatty acid) analysis

Phospholipid fatty acid data were obtained from NEON DP1.10104.001 for all available sites and dates using the neonUtilities R package (Lunch CK, 2020). Detailed data collection methods are available in (L. Stanish, 2020). Briefly, field-moist soils were sieved to 2mm or hand-picked to remove debris, before storing at -80° C for transport to an external lab for processing. Total phospholipid content was extracted from soils before discrete lipid molecules were identified using mass spectrometry and gas chromatography. Lipid peaks were compared to a reference standard with known concentrations of over 50 lipid compounds. Lipid amounts were then scaled by lipid extraction efficiency (Jackoway, 2020). The sum of all lipids attributable to fungal or bacterial lineages were used to calculate fungal:bacterial ratios (Lewe et al., 2021). Lipids chosen to represent each lineage and corresponding citations are listed in Table S1. Scripts to perform these calculations are hosted on Github (https://github.com/zoey-rw/SoilBiomassNEON).

#### ITS rRNA sequencing

Fungal taxonomic data were generated from ITS rRNA sequences distributed in NEON DP1.10109.001 (L. F. Stanish & Parnell, 2018) and processed according to details in (Werbin et al., 2024). To generate these data, soil cores were sampled at up to three time points per year at each NEON sampling plot (L. F. Stanish & Parnell, 2018). DNA was extracted from soil using the PowerSoil HTP Kit (Battelle Ecology, Inc, 2017; Sperling et al., n.d.)), then sequenced using Illumina MiSeq v3 2×300 base-pair paired end chemistry at Battelle Applied Genomics (Battelle Memorial Institute, 2018). Sequences were clustered into exact sequence variants (ESVs) using the dada2 pipeline before assigning taxonomy with the UNITE (Nilsson et al., 2019) and UTOPIA (Theil & Rifa, 2019) databases (Werbin et al., 2024).

### Mycorrhizal tree area

The basal area of mycorrhizal-associated tree species was calculated for seven NEON sites by (Lang et al., 2023). Briefly, the Vegetation Structure data product (DP1.10098.001) was used to determine biomass of live trees with either species or genus identifications during the most recent sampling event. These were then matched to the USDA PLANTS database (USDA NRCS & National Plant Data Team, 2021) to identify mycorrhizal associations. The area of arbuscular mycorrhizal (AM-) or ectomycorrhizal (ECM-) associated trees was then divided by total basal area to determine the proportion of each tree category.

## RESULTS

### Improved taxonomic classification of soil metagenomes compared to existing databases

SoilMicrobeDB classifies a higher percentage of reads to species from soil metagenomes than existing genome databases (Figure 2). Classification using SoilMicrobeDB was roughly double that of the PlusPF database (p < 0.001; Figure 2A), which was constructed from more than twice as many genomes. Initial read classification, using Kraken2 output without false-positive filtering, resulted in 41% reads classified with SoilMicrobeDB, which was significantly higher (p < 0.001) than initial read classification with GTDB (36% reads classified). After filtering to high-quality classifications at the domain level and normalizing by genome length (see Methods), 18% of reads were retained from the SoilMicrobeDB, compared to 15% from the GTDB (Figure 2B), which does not include fungi or novel soil MAGs.

**Figure 2.**
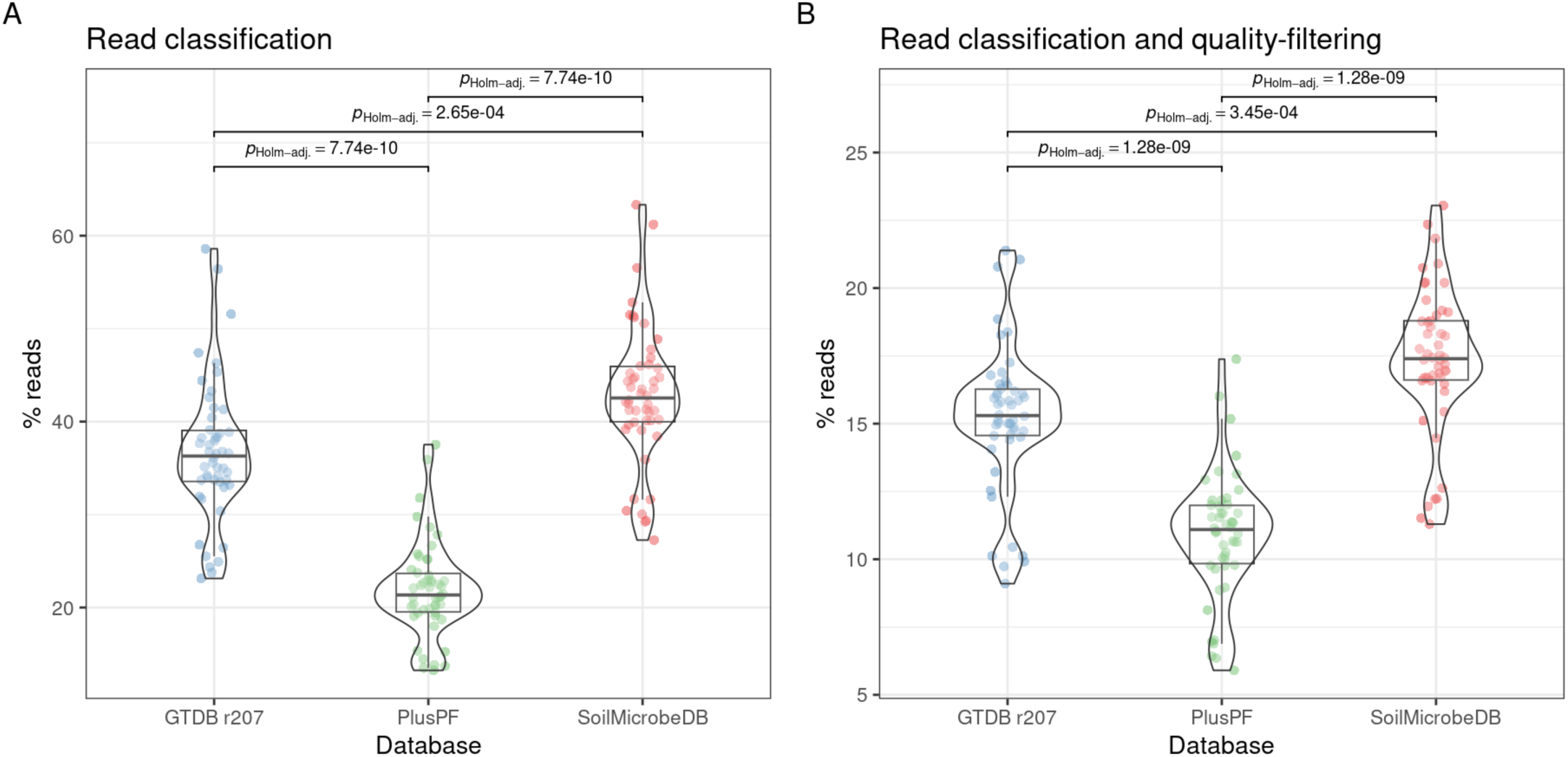
Metagenomic read classification of NEON soil samples at the domain level using three genome databases and filtering for high-quality classifications (using consistency of per-read kmer classifications) with Architeuthis. Brackens indicate significant Holm-adjusted p-values of Games-Howell test between groups.

Key lineages of fungi are the source of the improved classification through SoilMicrobeDB. Compared with the GTDB, which includes bacteria and archaea, the SoilMicrobeDB estimates a lower relative abundance of bacteria and a comparable relative abundance of archaea (Supplementary Figure 1). However, compared to PlusPF, which includes RefSeq fungi, the median estimated abundance of fungi doubled from 3% to 6%, due to increased classification of ectomycorrhizal fungal genera, such as *Cenococcum* and *Lactifluus*, which were represented in the Mycocosm fungal genome collection (Supplementary Figure 2).

### Mock community results validate the accuracy of SoilMicrobeDB

When mock communities of 200 species with known genome proportions were classified, the SoilMicrobeDB performed favorably compared to the other databases. The SoilMicrobeDB accurately recovered the abundances of fungi, bacteria, and archaea in the mock communities with minimal variation in accuracy across replicates of different sequencing depths (Figure 3C). The prokaryote-only GTDB overestimated most abundances and misclassified fungal reads to bacterial taxa, whereas the PlusPF database overestimated some species and underestimated other species (Supplementary Figure 3). We also found that higher proportions of fungi in the mock community were associated with less accurate estimates from PlusPF or GTDB (Figure 3C), while the sequencing depth of simulated samples did not affect the accuracy of any database.

**Figure 3.**
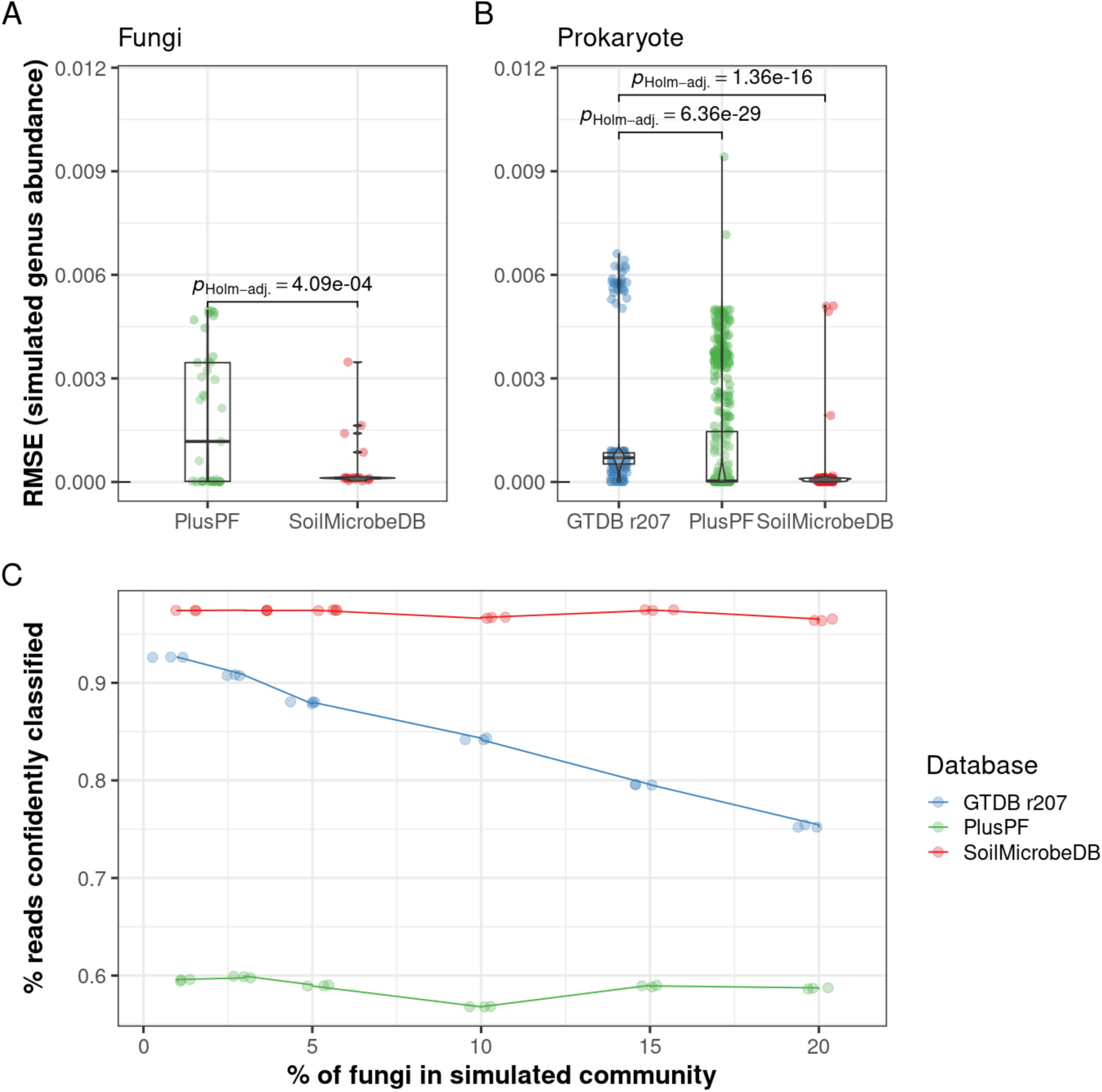
Performance of 3 databases against a simulated community of 200 species, 90% of which were prokaryote (bacteria or archaea) genera common to all databases, and 10% of which were fungi common to the PlusPF and SoilMicrobeDB. Root mean squared error (RMSE) of each taxon across simulated sequencing depths, where more accurate predictions are closer to zero, for **A)** fungi or **B)** prokaryotes. **C)** Percent reads classified for simulated communities with varying proportions of fungal genomes. The overlapping points are the different sequencing depths simulated (from .5m to 20M), which produced almost identical estimates.

### Metagenomic fungal-bacterial ratios and fungal relative abundances are consistent with biomass and rRNA-derived estimates

Consistent with our second hypothesis, the relative abundance of fungi in soil estimated from the SoilMicrobeDB correlated positively with fungal abundances estimated from other methods (Figure 4). The correlation was strongest for PLFA data (Figure 4A; R = .56, p < .0001), which closely reflects active biomass of fungi and bacteria but excludes archaea. The correlation with qPCR ratios was also positive (R = .19), but the relationship explained much less of the variation in fungal abundances (Figure 4B; R^2^ = .036, p < 0.0001).

**Figure 4.**
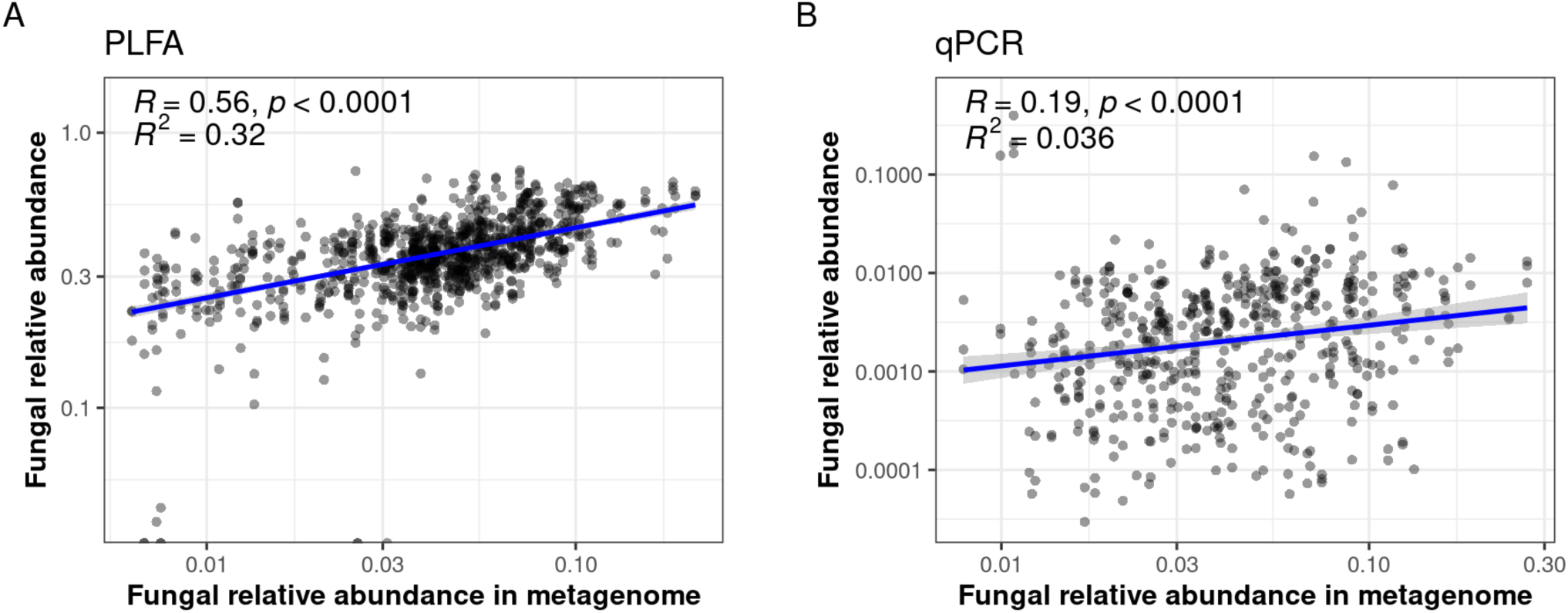
Relationships between relative abundance of fungi to bacteria in soil samples collected by the National Ecological Observatory Network, estimated using SoilMicrobeDB (x-axis) and **A)** PLFA (phospholipid fatty acid) abundance (y-axis) or **B)** quantitative PCR (y-axis). Black line represents 1:1 relationship between data types. **C)** Spearman correlation (R) between genus abundances estimated from ITS rRNA sequencing and metagenome sequencing for 100 most abundant fungal genera, grouped into functional guild membership. The 8 guild combinations with the most taxa are shown.

When the metagenomic-derived relative abundances of specific fungal lineages were compared with estimates from a widely used amplicon sequencing approach (amplification of internal transcribed spacer 1 or ITS1 rRNA (Thompson et al., 2018)), many genera showed agreement between approaches, including widespread ectomycorrhizal fungi (EMF) such as *Cenococcum*, *Russula*, and *Lactarius* (Figure 5A). Certain arbuscular mycorrhizal fungi (AMF) genera (e.g., *Glomus)*, which are known to be under-estimated by ITS sequencing (Lekberg et al., 2018), had weaker relationships with metagenome estimates (Figure 5A). However, metagenomic estimates of AMF were strongly correlated with AMF-associated trees (Figure 5B; R = .52, R^2^ = .27, p < 0.0001).

**Figure 5.**
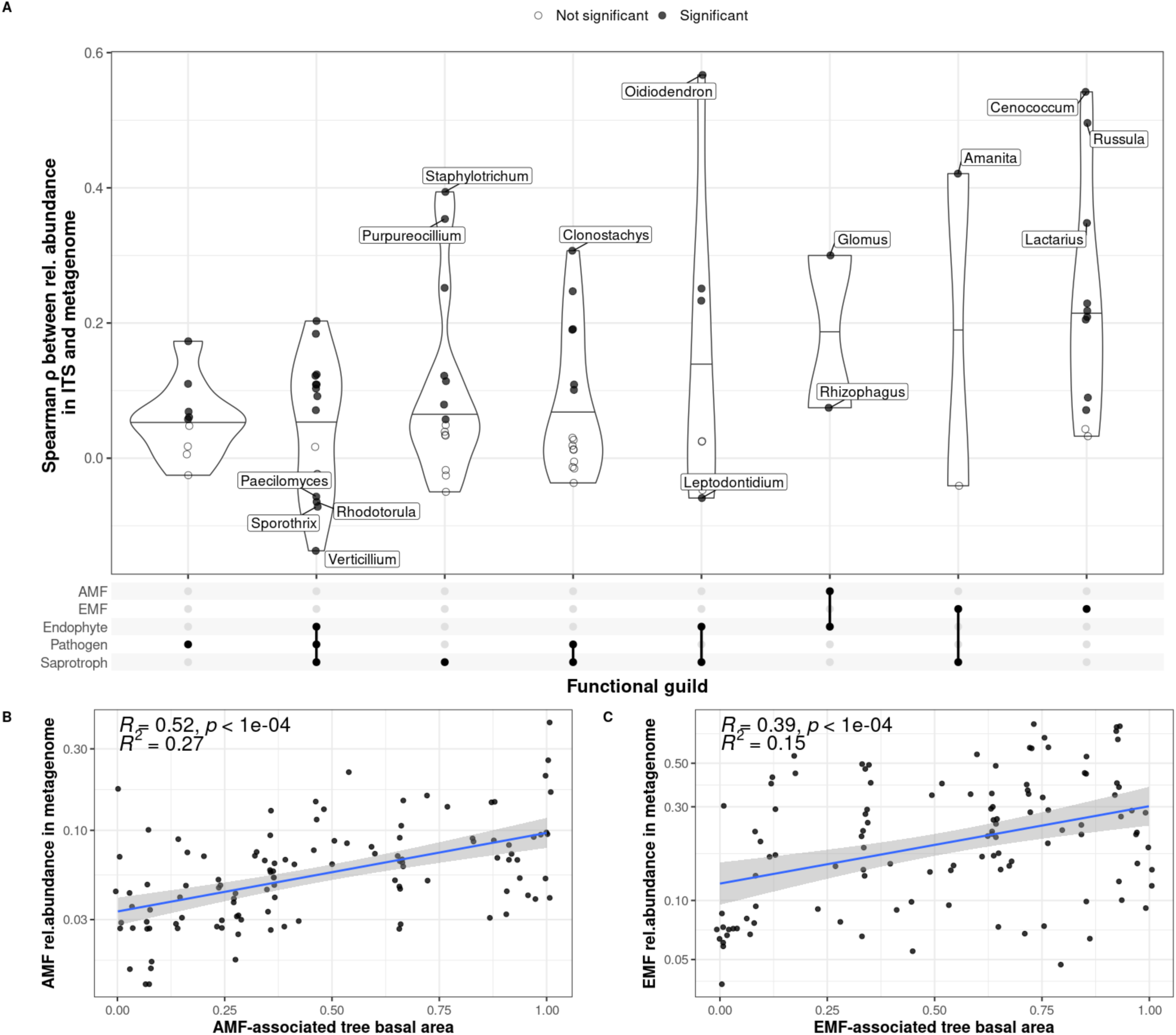
Relationship between metagenome-estimated mycorrhizal fungi relative abundance and mycorrhizal-associated tree basal area. **A)** Y-axis shows arbuscular mycorrhizal fungi (AMF) abundance, expressed as a proportion of all fungal abundances. **B)**: Y-axis shows ectomycorrhizal fungi (EMF) abundance, expressed as a proportion of all fungal abundances within ectomycorrhizal genera (Tedersoo & Smith 2013). AM or EM tree basal area was calculated and reported in Lang et al. 2023.

### Biomes differ in the dominance of uncultured organisms

In support of our third hypothesis, we found variation between biomes in the proportion of soil DNA that matched to uncultured organisms (MAGs). The median abundance of MAGs ranged from 20% in agricultural biomes, such as croplands and pastures, to over 35% in boreal (Alaskan) shrublands, wetlands, and herbaceous biomes (Figure 6A). We also found that the samples dominated by uncultured organisms tended to have lower classification rates (Figure 6B). This relationship between classifiability and culturability is present across samples, but the relationship explains less variability in Alaska samples, which had higher overall abundances of uncultured organisms (Figure 6B).

**Figure 6.**
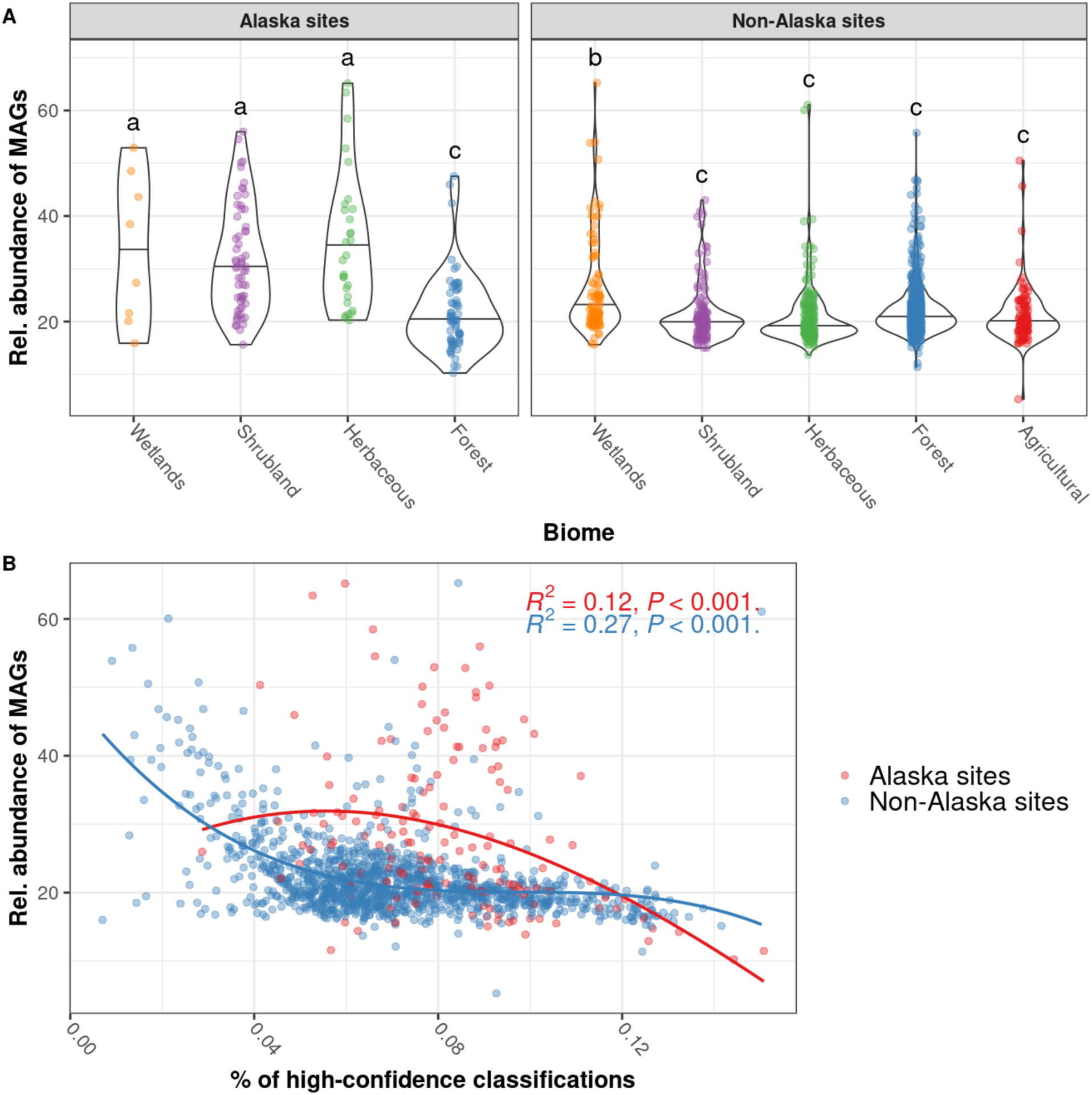
SoilMicrobeDB classification of metagenomes sequenced by the National Ecological Observatory Network (NEON). **A)** Relative abundance of MAGs across biome types. Biome type was converted from each sample’s NLCD class using the NLCD2011 Legend (Multi-Resolution Land Characteristics (MRLC) consortium, 2011). Letters indicate significantly different groups (Tukey HSD after ANOVA with p < .001). **B)** Relative abundance of MAGs (y-axis) and percent of high-confidence classifications (x-axis). High-confidence classifications are the Kraken2-assigned reads passing the kmer-consistency quality filter applied by Architeuthis. Lines indicate polynomial regression implemented in the ggpmisc R package (Aphalo, 2016), with the formula specified as y ∼ poly(x, 3, raw = TRUE), fitted separately to Alaska sites (red) and non-Alaska sites (blue).

### Interactive web application for data accessibility

An interactive web application was developed to allow data exploration and download of output files (Figure 7). By linking to complementary metadata on various microbes, such as culture conditions and the availability of *in silico* modeling tools, this portal reports the current state of research on each organism and serves as a resource for downstream applications such as biogeography, ecological, and predictive modeling studies. Each microbe within the SoilMicrobeDB falls into one of the following four categories:

- *Uncultured*: Metagenome-assembled genome (MAG) or single-cell assembled genome (SAG): These microorganisms have no reported cultures and are primarily studied using genomic approaches. This currently includes 49% of genomes within the database.
- *Isolate genome:* These genomes were derived from an isolate and consequently, the organism is assumed to have survived in culture. This currently includes 51% of genomes within the database.
- *Well-characterized:* These microorganisms have been cultured and have at least one experimentally-validated genome-scale model (GEM) that can be used to simulate metabolism and physiology under both observed and novel conditions. This includes <100 taxa.
- *Model organism*: These microorganisms have been studied by multiple research teams and avenues, ideally with ’omic and experimental data integrated as a Digital Microbe (Veseli et al., 2024). This includes <20 taxa.

**Figure 7.**
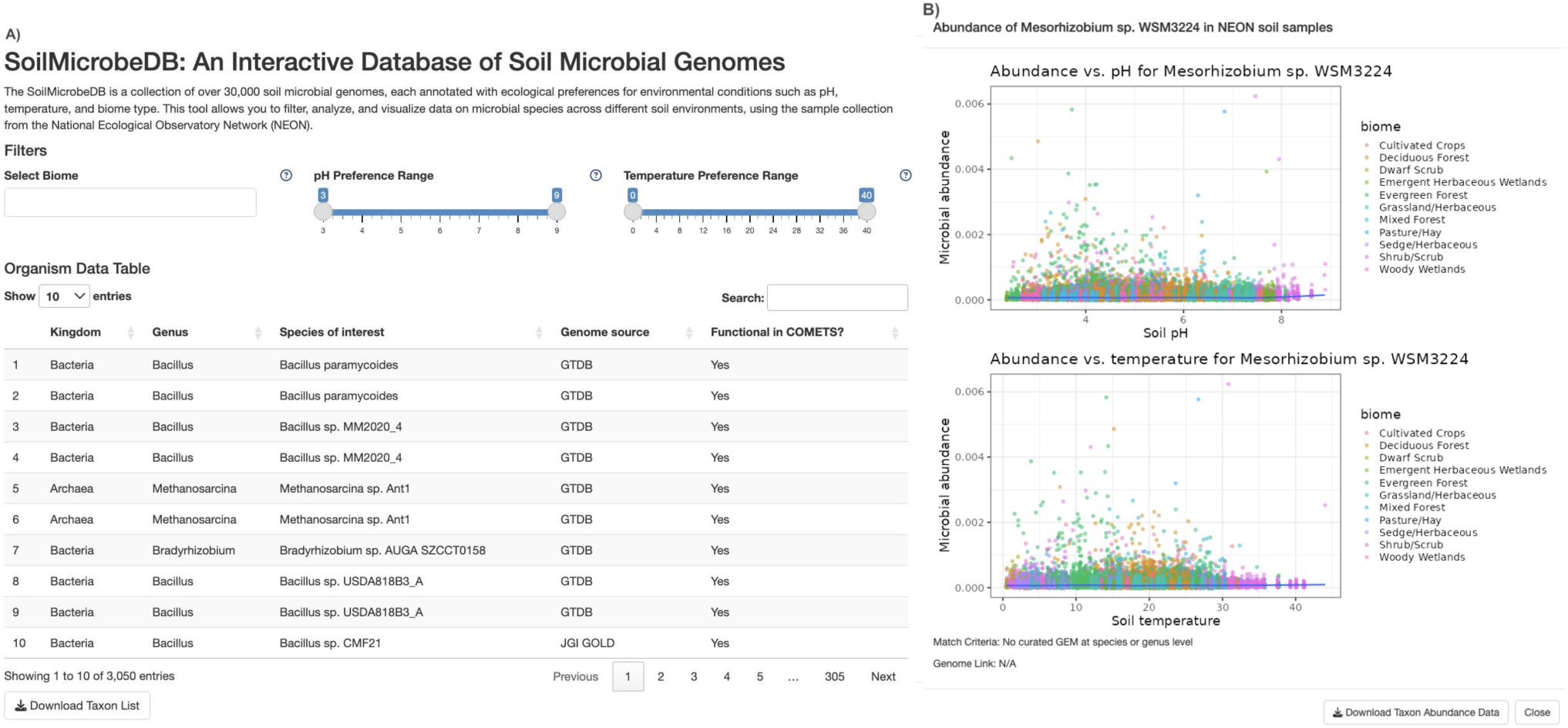
Interactive web application developed to distribute the results from mapping metagenomes against the SoilMicrobeDB. A) home page displaying all taxa within the SoilMicrobeDB, the source of each taxon’s genome, and whether or not the taxon has a genome-scale metabolic model (GEM) available. B) Selection of an individual taxon displays the abundance plotted against pH and temperature, associated publications with GEMs, and an option to download the data used for visualization. https://zoeywerbin.shinyapps.io/soil_microbe_db/.

To illustrate how this database can facilitate identification of microbes that could be promising candidates for bioprospecting, bioremediation, or terraforming, we assign ecological “preferences” to each microbe using environmental data (pH and temperature) associated with each NEON soil sample. These preferences were assigned based on the peak of a generalized additive model fit to observed abundances across environmental gradients (Tomal & Ciborowski, 2020; Trexler & Travis, 1993). Most preferences were clustered at the extremes of the temperature or pH range (Supplementary Figure 6). Each microbe can be inspected to view its abundance trends across pH and temperature ranges (Figure 7B). Abundances can then be downloaded for each microbe or for all microbes within the SoilMicrobeDB. The application can be accessed at https://zoeywerbin.shinyapps.io/soil_microbe_db/.

## DISCUSSION

Large-scale sequencing of environmental microbiomes has revealed that most soil organisms remain uncharacterized: most soil DNA does not map to any known domain of life (Anthony et al., 2024), and that most soil organisms cannot be cultured in a laboratory setting (Bartelme et al., 2020; Nesme et al., 2016). To address this issue, we developed SoilMicrobeDB, a database of soil-associated reference genomes for classifying metagenomic sequences. Mapping metagenomic reads to SoilMicrobeDB increased the taxonomic classification of soil samples by over 20% (Figure 2), with improvements attributed to the inclusion of fungal genomes and recently-published metagenome-assembled genomes (MAGs). When evaluating accuracy of taxonomic assignments using simulated data, SoilMicrobeDB outperformed comparison databases (Figure 3). Metagenomic results are consistent with biomass and rRNA-derived estimates of fungal abundances in soil (Figure 4), especially for ectomycorrhizal fungi (Figure 5). When comparing soil communities across biomes, the incorporation of MAGs increased the identifiable composition of boreal and tundra biomes in Alaska, which were dominated by yet-uncultured organisms (Figure 6).

A key advancement in this work is the demonstration that short-read metagenomes contain reliable signals of fungal abundance. Fungal abundance, often reported as a fungal-bacterial ratio, is widely used as a simple proxies for soil health within agriculture (Malik et al., 2016; Strickland & Rousk, 2010; Wang et al., 2019; Yu et al., 2022) and its correspondence with metagenomic-derived ratios suggests that vast amounts of already-available metagenomic sequence data could provide insights into spatiotemporal trends in soil health. We found that metagenomic abundances of fungi determined with SoilMicrobeDB were highly correlated with estimates from phospholipid fatty acid data (PLFA), but only weakly correlated with quantitative PCR (qPCR) data. PLFA is generally considered a better representation of living microbial biomass, whereas qPCR may be more sensitive to relic DNA and fungal phylogenetic biases (Siles et al., 2024). Given these differences, we conclude that the metagenome-derived abundances are reflecting cellular biomass indicators of the fungal community rather than DNA abundances. Another line of evidence supporting this conclusion is the correspondence between mycorrhizal tree status (i.e. the fraction of trees classified as arbuscular or ectomycorrhizal tree) and mycorrhizal abundances within the metagenome (Figure 5B-C). The improved detection of arbuscular and ectomycorrhizal fungi with SoilMicrobeDB suggests a new avenue for benchmarking the current distribution of fungal functional groups within soils, including those that play key roles within terrestrial ecosystems (Baldrian, 2019; Read & Perez-Moreno, 2003; Talbot et al., 2008).

Genomes are rapidly being assembled and published as MAG collections, but the ecological roles of taxa represented by these genomes remain unknown. By connecting MAGs with their observed environmental preferences, the SoilMicrobeDB will encourage the use of MAGs as a resource for efforts such as bioprospecting (Eren & Delmont, 2024). Our analysis also revealed a relationship between classifiability and culturability, suggesting that samples with high abundance of uncultured organisms and low classifiability likely represent sources of relative genetic novelty. Samples from Alaskan wetlands, shrubland, and herbaceous biomes had particularly high abundances of uncultured taxa (Figure 6). This may be due to climate conditions that are particularly extreme, or to novel microbes released from thawing permafrost (Ernakovich et al., 2022). NEON sites within Alaska are thawing at different speeds (e.g. a high rate of permafrost thaw has been observed at Healy Valley (Mulligan, Dennis, 2017)), so the ability to track microbial abundances across this thaw gradient is a valuable opportunity to improve understanding of Arctic soil microbial processes. Given the high abundances of uncultured taxa, we recommend that future studies use tools like GenomeSPOT for developing culture strategies or predicting phenotypes (Barnum et al., 2024) to improve our understanding of the ecological significance of these taxa.

The proportion of unclassified reads within soils remains much higher than other habitat types (Anthony et al., 2024). An unknown portion (up to 40%) of the DNA in a soil sample could be relic DNA from dead organisms (Carini et al., 2016). The effect of relic DNA is unknown, however these sequences should theoretically be classifiable. Instead, low classification of soil DNA may occur because, compared to human-associated or engineered ecosystems, a very large portion of soil functional genes remain truly uncharacterized (Pavlopoulos et al., 2023). Many genes within metagenomes may belong to this class of functional genes, rather than ribosomal genes that have been often targeted for identification. However, analyses of unidentifiable DNA suggest that entire lineages are still unknown to science: gene family prediction of unique traits within unidentifiable environmental sequences suggests that there are at least 18 undiscovered prokaryotic phyla (Rodríguez Del Río et al., 2024). Alternative explanations may be the prevalence of repetitive or non-coding sequences that are not complex enough to find a match using 150bp reads ((Clarke et al., 2019)). If this is the case, then longer-read sequencing technologies such as Oxford Nanopore and Pacific Biosciences will likely help estimate the percentage of environmental DNA made up of non-coding sequences. The landscape of available fungal genomes is likely to change dramatically over the next decade. About half of genomes within the Mycocosm collection - over 1000 - are currently in a “restricted” category requiring the permission of each individual genome submitter to be used in an analysis, but this category expires after 2 years, after which point the genomes will be freely accessible according to the JGI data policy. Additionally, tools such as EukCC (Saary et al., 2020) have been developed to assemble eukaryotic genomes directly from metagenomes, which will further increase the availability of fungal soil genomes. Approximately 23% of fungal reads could be confidently classified to the rank of phylum but not to genus, but expanding the fungal diversity of the SoilMicrobeDB should increase the proportion of DNA that can be identified at finer ranks.

Shotgun metagenomics has remained a less accessible method for ecologists, compared to marker gene sequencing, due to higher costs of sequencing and the complexity of raw metagenomic data. We aimed to reduce these barriers to entry by providing an interactive portal that presents the abundances of each of 52,193 organisms within the SoilMicrobeDB across key environmental gradients. These abundance data can be filtered and downloaded for subsequent analyses, e.g. identifying potential extremophile strains for astrobiological applications. For instance, filtering by low-temperature preferences highlights species within the genus *Deinococcus*, which is a candidate organism for survival in Martian soils (Horne et al., 2022; Vora et al., 2024). The entire SoilMicrobeDB will be distributed free of charge until 2027 via the Amazon Web Services (AWS) Public Dataset program. Downloading the database allows for custom modification with additional genomes using the software Struo2 (Youngblut & Ley, 2021). This flexibility is critical for ensuring that as new genomes are assembled, the database can expand in coverage.

## Author Contributions

Conceptualization: ZW, MF; Data Curation: ZW; Formal Analysis: ZW; Funding Acquisition: JB, MF, DS; Investigation: ZW; Methodology: ZW, ID; Project Administration: MF; Resources: JB, MF; Software: ZW, ID; Supervision: JB, MF, DS; Visualization: ZW; Writing – Original Draft: ZW; Writing – Review & Editing: JB, ID, MF, DM, DS, WEA

## Data Availability

All raw data are publicly available. Processed data is available through the interactive web portal at [URL] and archived on Zenodo at time of publication.

## Code Availability

https://github.com/zoey-rw/SoilMicrobeDB

## Funding

ZW - National Science Foundation Graduate Research Fellowship Program #2020285775

JB, ZW - National Science Foundation (NSF) Macrosystems Biology grant #1638577

JB - BU Bioinformatics Research and Interdisciplinary Training Experience (BRITE) NSF-REU program #1949968

MRF, ZW - Rafik B. Hariri Institute for Computing and Computational Science & Engineering Focused Research Program #2023-07-002: “First Trip to Mars: How to Pack Light”

## Supplementary Materials

**Supplementary Figure 1.**
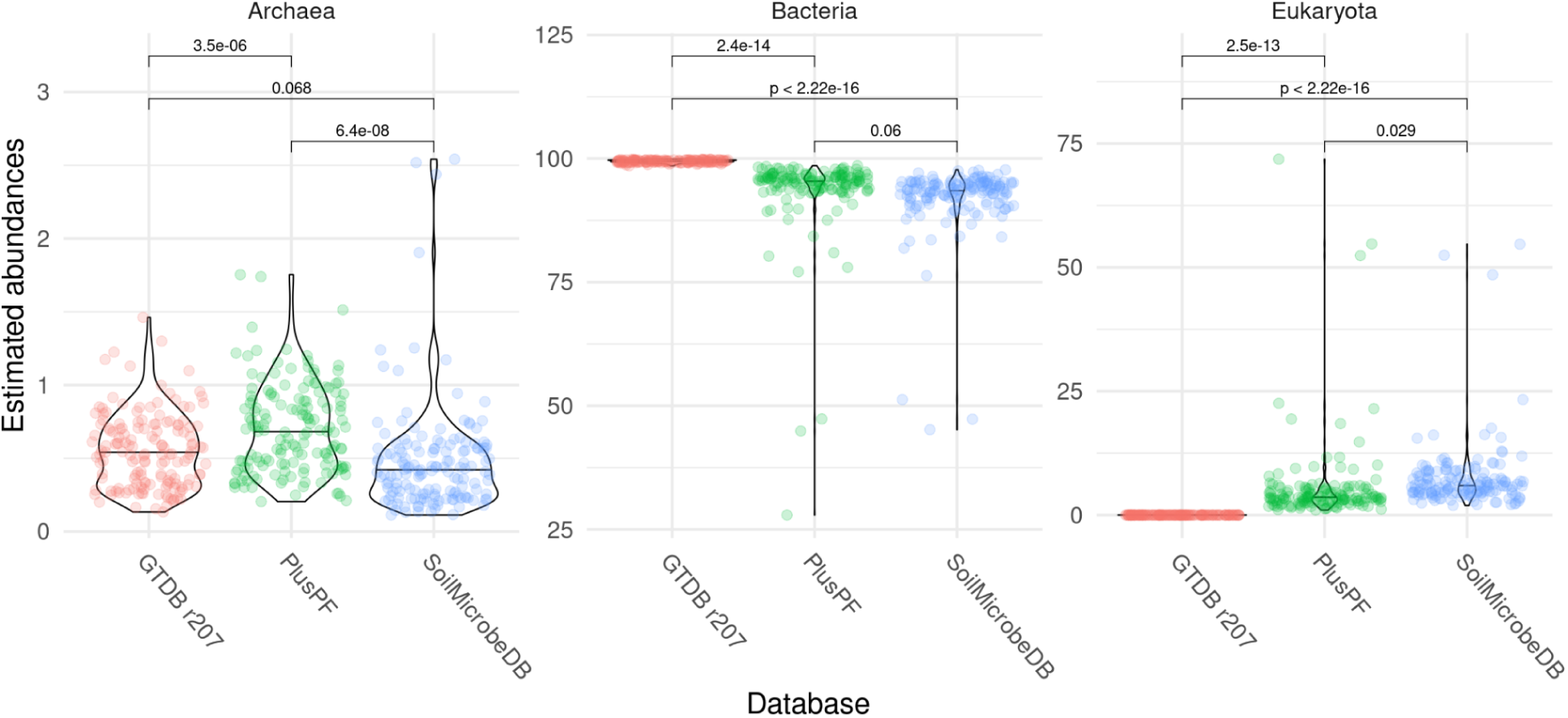
Estimated relative abundance of microbial domains at NEON sampling sites. Reads were classified using three Kraken2 databases (SoilMicrobeDB, PlusPF, and GTDB r207), followed by kmer filtering with Architeuthis, and relative abundance estimation using Bracken. Brackets show p-values for T-tests between databases.

**Supplementary Figure 2.**
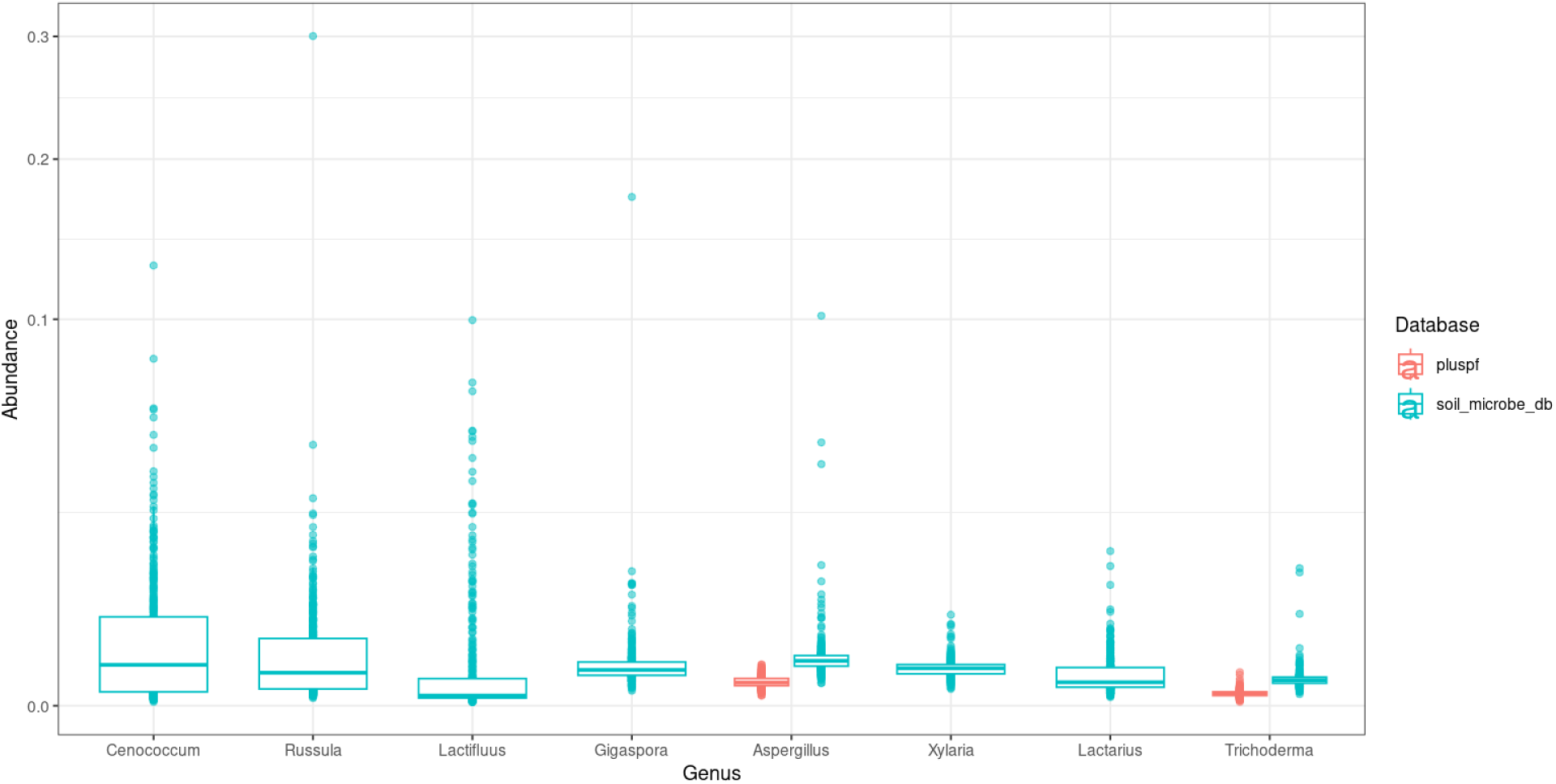
Abundances of widespread soil fungal genera. Reads were classified using two Kraken2 databases (SoilMicrobeDB and PlusPF), followed by kmer filtering with Architeuthis, and relative abundance estimation using Bracken.

**Supplementary Figure 3.**
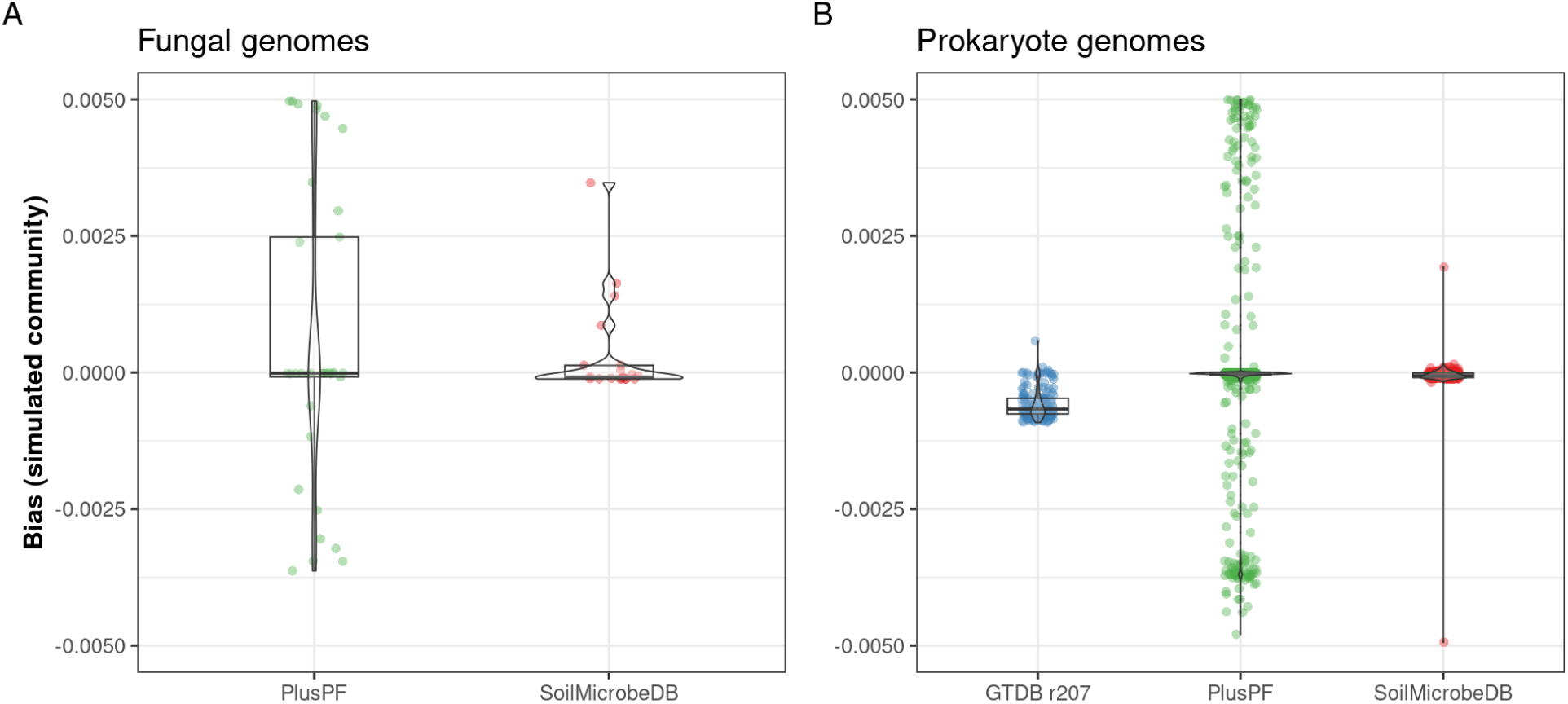
Mean bias of 3 database relative abundance estimates for taxa within a simulated community of 200 species, 10% of which were fungi common to the PlusPF and SoilMicrobeDB (panel A) and 90% of which were prokaryote (bacteria or archaea) genera common to all databases (panel B). Less biased predictions are nearer to zero (indicated by the horizontal dotted line).

**Supplementary Figure 4.**
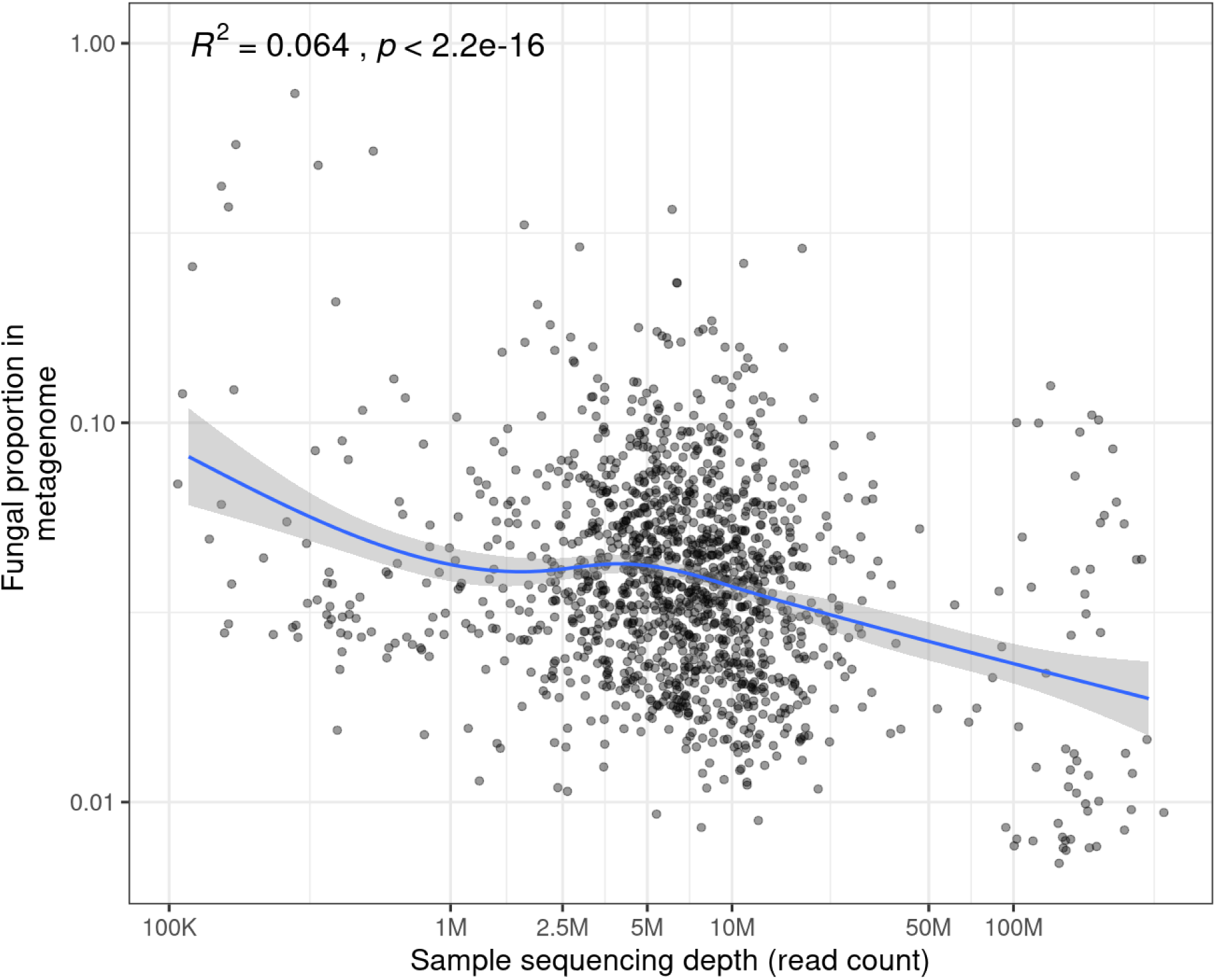
Negative relationships between sequencing depth (x-axis, log scale) and the relative abundance of fungi (y-axis, log scale) detected in NEON soil samples, using SoilMicrobeDB. Plotted line and values show generalized additive model (GAM) regression.

**Supplementary Figure 5.**
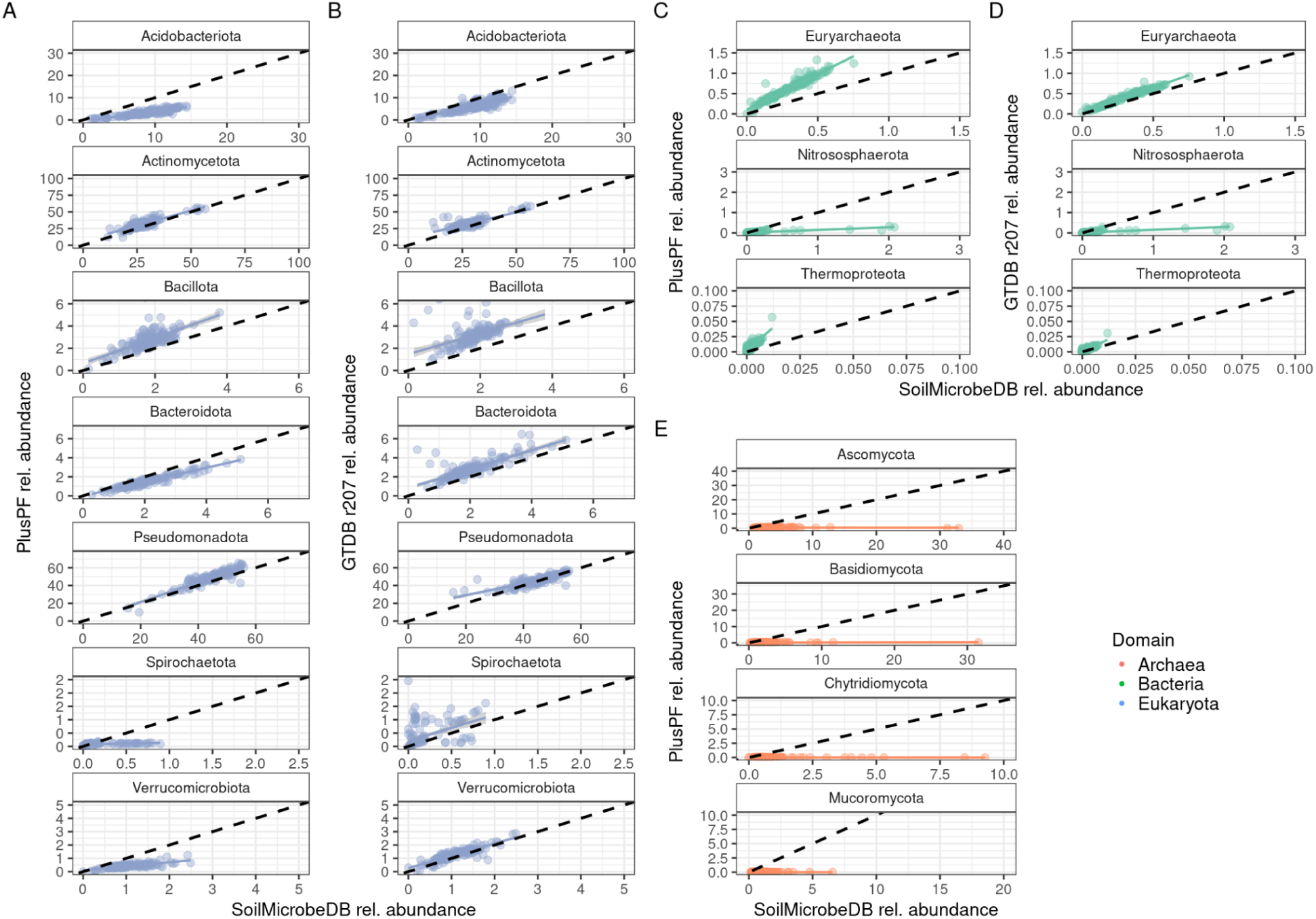
Relative abundances of soil microbial phyla estimated by different databases. SoilMicrobeDB (x-axis) contains soil-associated bacteria and archaea, and fungi. PlusPF database (y-axis for panels A, C, and E) includes Refseq archaea, bacteria, viral, plasmid, human, UniVec_Core. GTDB r207 database (y-axis for panels B and D) includes archaea and bacteria. Black dashed line represents 1:1 line. After classifying sequences with Kraken2, and filtering false-positives using Architeuthis, relative abundances were estimated by Bracken using normalization by kmer coverage and genome size.

**Supplementary Figure 6.**
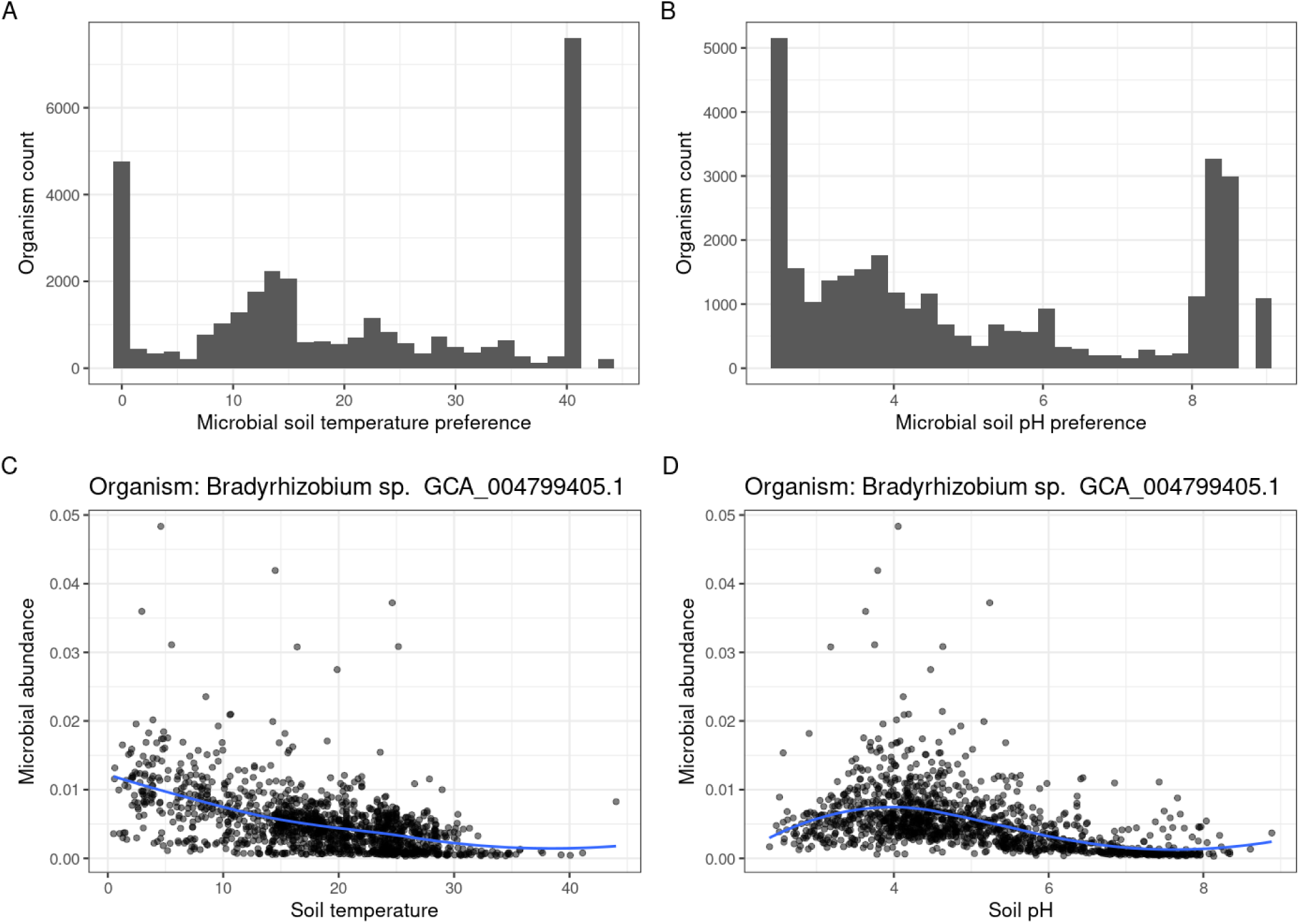
Observed A) soil temperature or B) soil pH preferences of microbes in SoilMicrobeDB (n = 26258 organisms detected) mapped to short read metagenomes from NEON (n = 1385 soil samples). Preferences were assigned based on the peak of a LOESS model fit in R using the ggplot2 function stat_smooth(method=“loess”). Example for one organism shown in panels C) and D), which show the LOESS fit (blue line) for abundances across C) soil temperature or D) soil pH.

**Supplementary Table 1:** Data sources for individual phospholipid fatty acid (PLFA) classifications used to estimate fungal or bacterial abundances from NEON soil samples, published as NEON DP1.10104.001. (Docherty et al., 2015; Giray et al., 2024; Joergensen, 2022; Lewe et al., 2021).

| Variable name | Description | Target Lipid | Taxonomic group | Broad taxonomic group |
| --- | --- | --- | --- | --- |
| cis18To2n912ScaledConcentration | cis,cis-9,12-octadecadienoic acid, or linoleic acid, or 18To2-omega-6, methyl ester, cis18:2n9-12 | 18:2 ω6,9c | general fungi minus AMF | Fungi |
| c16To1Cis11ScaledConcentration | methyl cis-11-hexadecenoate, 16:1 cis 11 | 16:1 w5c (11-Hexadecenoic acid) | AMF | Fungi |
| cis16To1n9ScaledConcentration | methyl hexadecenoic acid, or palmitoleic acid, methyl ester, c16:1n9 | 16:1 w7c | Gram-negative Bacteria | Bacteria |
| i15To0ScaledConcentration | 13-methyltetradecanoic acid methyl ester, i15:0 | i15:0 | Gram-positive Bacteria | Bacteria |
| c17To0AnteisoScaledConcentration | methyl 14-methyl-pentadecanoate, 17:0 anteiso | a17:0 | Gram-positive Bacteria | Bacteria |
| lipid10Methyl18To0ScaledConcentration | tuberculostearic acid methyl ester, 10 Me-18:0 | 10 Me-18:0 | Gram-positive Bacteria | Bacteria |
| lipid10Methyl16To0ScaledConcentration | methyl 10-methyl-hexadecanoate, 10 Me-16:0 | 10 Me-16:0 | Gram-positive Bacteria | Bacteria |
| cyclo17To0ScaledConcentration | C17To0 fatty acid methyl cis-9,10-methylenehexadecanoate, cyclo17:0 | cyclopropyl fatty acids | Gram-negative Bacteria | Bacteria |
| cyclo19To0ScaledConcentration | C19To0 fatty acid methyl cis-9,10-methyleneoctadecanoate, cyclo19:0 | cyclopropyl fatty acids | Gram-negative Bacteria | Bacteria |
| c18To3n6ScaledConcentration | 6,9,12-octadecatrienoic acid, or gamma-linolenic acid, methyl ester, c18:3n6, | 18:3ω6,9,12c |  | Fungi |
| c18To3n3ScaledConcentration | cis,cis,cis-9,12,15-octadecatrienoic acid, or alpha-linolenic acid, methyl ester, c18:3n3 | 18:3ω3,6,9 | saprotrophic and EMF | Fungi |
| c14To0ScaledConcentration | tetradecanoic acid, or myristic acid, methyl ester, c14:0 | 14:0 |  | Microbial |
| c15To0ScaledConcentration | pentadecanoic acid methyl ester, c15:0 | 15:0 |  | Microbial |
| c16To0ScaledConcentration | hexadecanoic acid, or palmitic acid, methyl ester, c16:0 | 16:0 |  | Microbial |
| c17To0ScaledConcentration | heptadecanoic acid methyl ester, c17:0 | 17:0 |  | Microbial |
| c18To0ScaledConcentration | octadecanoic acid, or stearic acid, methyl ester, c18:0 | 18:0 |  | Microbial |
| aC15To0ScaledConcentration | 12-methyltetradecanoic acid methyl ester, a-c15:0, | a15:0 | Gram-positive Bacteria | Bacteria |
| i16To0ScaledConcentration | methyl 14-methylpentadecanoate, i16:0 | i16:0,<br>14-methylpentadecanoic acid | Gram-positive Bacteria | Bacteria |
| i17To0ScaledConcentration | methyl 15-methylhexadecanoate, i17:0 | i17:0,<br>15-methylhexadecanoic acid | Gram-positive Bacteria | Bacteria |
| lipid10Methyl17To0ScaledConcentration | heptadecanoic acid methyl ester, 10 Me-17:0 | 10Me17:0 | Gram-positive Bacteria | Bacteria |
| lipid2OH10To0ScaledConcentration | methyl 2-hydroxydecanoate, 2OH10:0 | 2OH10:0 | Gram-negative Bacteria | Bacteria |
| lipid2OH12To0ScaledConcentration | methyl 2-hydroxydodecanoate, 2OH12:0 | 2OH12:0 | Gram-negative Bacteria | Bacteria |
| lipid2OH14To0ScaledConcentration | methyl 2-hydroxytetradecanoate, 2OH14:0 | 2OH14:0 | Gram-negative Bacteria | Bacteria |
| lipid2OH16To0ScaledConcentration | methyl 2-hydroxyhexadecanoate 2OH16:0 | 2OH16:0 | Gram-negative Bacteria | Bacteria |
| lipid3OH12To0ScaledConcentration | methyl 3-hydroxydodecanoate, 3OH12:0 | 3OH12:0 | Gram-negative Bacteria | Bacteria |
| lipid3OH14To0ScaledConcentration | Concentration of methyl 3-hydroxytetradecanoate, 3OH14:0 | 3OH14:0 | Gram-negative Bacteria | Bacteria |
| cis18To1n9Scaled<br>Concentration | cis-9-octadecenoic acid, or oleic<br>acid, methyl ester, cis18:1n9, | C18:1ω9 | saprotrophic<br>fungi | Fungi |

